# Nanoscale spatio-temporal diffusion modes measured by simultaneous confocal and STED imaging

**DOI:** 10.1101/287680

**Authors:** Falk Schneider, Dominic Waithe, Silvia Galiani, Jorge Bernadino de la Serna, Erdinc Sezgin, Christian Eggeling

## Abstract

The diffusion dynamics in the cellular plasma membrane provides crucial insights into the molecular interactions, organization and bioactivity. Fluorescence correlation spectroscopy combined with super-resolution stimulated emission depletion nanoscopy (STED-FCS) measures such dynamics with high spatial and temporal resolution and reveals nanoscale diffusion characteristics by measuring the molecular diffusion in conventional confocal mode and super-resolved STED mode sequentially. However, to directly link the spatial and the temporal information, a method that simultaneously measures the diffusion in confocal and STED modes is needed. Here, to overcome this problem, we establish an advanced STED-FCS measurement method; line interleaved excitation scanning STED-FCS (LIESS-FCS) which discloses the molecular diffusion modes at different spatial positions with a single measurement. It relies on fast beam-scanning along a line with alternating laser illumination that yields, for each pixel, the apparent diffusion coefficients for two different observation spot sizes (conventional confocal and super-resolved STED). We demonstrate the potential of the LIESS-FCS approach with simulations and experiments on lipid diffusion in model and live cell plasma membranes. We also apply LIESS-FCS to investigate the spatio-temporal organization of GPI-anchored proteins in the plasma membrane of live cells which interestingly show multiple diffusion modes at different spatial positions.

## Introduction

Lateral heterogeneity in plasma membrane organization is known to modulate cellular functionalities in a wide range of biological processes^1,2^. This heterogeneity and the underlying structures or molecular interaction dynamics can be probed through investigation of molecular diffusion characteristics in the plasma membrane over space and time ^3,4^. A widely employed approach to explore molecular diffusion in the plane of the cellular plasma membrane is fluorescence correlation spectroscopy (FCS). FCS is usually employed to determine the average transit times (τ_D_) of molecules through a confocal observation volume to obtain the diffusion coefficients (D), revealing changes in molecular diffusion due to, for example, changes in membrane viscosity or molecular interactions^5^. Additionally, non-Brownian hindered diffusion caused by molecular interactions and confinements has been studied using FCS^6^. Later, molecular diffusion modes (not only the overall velocity of the molecules but also the diffusion characteristics) in the plasma membrane were measured. For these measurements, FCS was employed at different length scales allowing the determination of transit times τ_D_ for observation spots (*d*) of varying sizes (ranging from ~200 nm to >µm)^7^. By plotting the dependence of τ_D_ on *d* (τ_D_(*d*)), such spot-variation FCS (svFCS) measurements were used to distinguish different diffusion modes such as free (Brownian) diffusion, transient trapping in slow moving or immobilized entities (trapped diffusion), or compartmentalized (hop) diffusion^8^. Unfortunately, parameters such as trapping times or sizes of the trapping sites could only be extrapolated (even in the case of more advanced camera-based approaches)^9,10^, since the relevant molecular scales are below the diffraction-limited spatial resolution of these techniques. The remedy to this was to make direct FCS recordings with sub-diffraction sized observation spots, as created by near-field illumination (necessitating the close proximity to nanostructured surfaces or apertures)^11,12^ or through super-resolution far-field STED microscopy^13,14^. Subsequently, STED-FCS diffusion modes were extracted from τ_D_(*d*) (or D(*d*)) dependencies ranging from diffraction-limited d~240nm down to d<50nm). To thoroughly understand the spatial heterogeneity and related spatial diffusion modes, FCS needs to be recorded simultaneously at various points, as done for scanning-FCS where multiple FCS data are recorded for each pixel along a quickly scanned line^15^. Consequently, scanning STED-FCS (sSTED-FCS) recordings for fluorescent lipid analogues in the plasma membrane of living cells revealed distinct transient sites of slowed-down diffusion that extended over <80nm^16^. sSTED-FCS so far has not been used to extract diffusion modes in these sites, since values of τ_D_ (or D) could only be determined for one observation spot diameter *d* at a time.

The sequential measurements (first confocal and then STED) are not optimal due to the cellular movements or fast molecular dynamics. This fundamental limitation of using only one diameter spot at a time restricts the application of STED-FCS measurements. The time needed in between confocal and STED measurements makes it impossible to gain the nanoscale spatio-temporal diffusion modes for the selected area. The only way to overcome this is scanning through a line with confocal and STED simultaneously. In this paper, we show an approach allowing (quasi-)simultaneous extraction of STED-FCS data for different *d* as previously achieved in single-point FCS which lacked the spatial aspect^17,18^. We present here line interleaved excitation scanning STED-FCS (LIESS-FCS), which by fast beam-scanning along a line with alternating laser illumination provides, for each pixel, apparent diffusion coefficients for two different observation spot sizes, one corresponding to the diffraction-limited confocal and the other to super-resolved STED. We validate our LIESS-FCS approach with simulations and employed it to investigate nanoscale molecular diffusion modes in the plasma membrane of live cells. We observed various diffusion modes for different lipid species and interestingly a combination of different diffusion characteristics for GPI-anchored proteins.

## Results

The basic principles of scanning STED-FCS (sSTED-FCS) and line interleaved excitation scanning STED-FCS (LIESS-FCS) are depicted in Fig. 1A and 1B. In sSTED-FCS, either the larger confocal (*d*_conf_~240nm) or smaller STED (*d*_STED_<<200nm) observation spot quickly and multiple times scans over the sample through a line (or a circle), creating temporal intensity data. Then, the correlation function is applied to the intensity trace at each pixel along the line, generating the autocorrelation functions summarised in space in the so-called correlation carpets (in either confocal or STED Fig. 1C, D). In sSTED-FCS, usually, values of an apparent diffusion coefficient for confocal, D_conf_ = D(*d*_conf_), and STED recordings, D_STED_ = D(*d*_STED_), can only be determined subsequently, not simultaneously. Therefore, D_conf_ and D_STED_ cannot be paired to determine spatially resolved D(*d*) dependencies since diffusion characteristics may have changed at the individual pixels in-between confocal and STED recordings. This means in conventional sSTED-FCS, cellular movements, variations in the plasma membrane topology, or any other heterogeneity in the plasma membrane display a significant challenge for subsequently recorded confocal and STED-FCS data. In contrast, in LIESS-FCS, the confocal and STED-based observation spots are scanned in an alternating manner (line-by-line basis), creating intensity and correlation carpets and thus values of D_conf_ and D_STED_ for each pixel quasi-simultaneously within the same measurement. This now allows relating values of D_conf_ and D_STED_ by calculating the ratio D_rat_ = D_STED_/D_conf_ for each pixel. Values of D_rat_ give unique information on the nanoscale diffusion characteristics, since they vary for different nanoscale diffusion modes as detailed before^19,20^: D_rat_ = 1 for free, D_rat_ < 1 for trapping and D_rat_> 1 for hop (or compartmentalized) diffusion.

**Figure 1:**
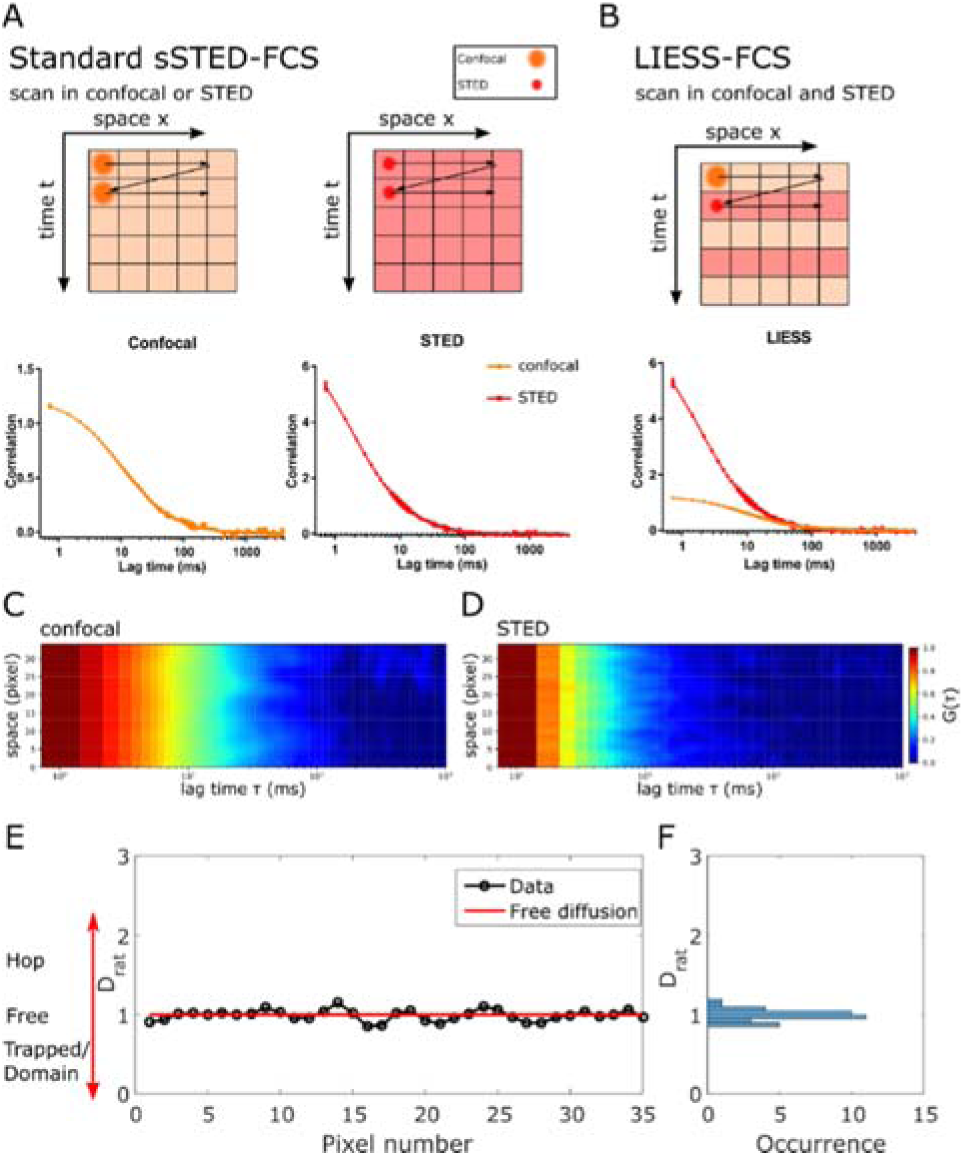
Principle of LIESS-FCS: **A)** sSTED-FCS data are usually generated from rapidly scanning with a diffraction-limited confocal (orange) and super-resolved STED (red) spot several times (time t axis) along a line (spatial x axis) yielding intensity trace for each pixel along the line which is then correlated to generate the final FCS data (Correlation data G(τ) against correlation lag time τ) in confocal and STED separately (bottom plots) **B)** In LIESS-FCS, confocal and super-resolved STED-FCS data are generated simultaneously by alternating in-between subsequent lines confocal and STED modes. Arrows: movement of the beam scanner. **C, D)** Representative correlation carpets in **C)** confocal and **D)** STED for simulated data of free diffusion (measurement time 40 s, *d*_STED_ = 100 nm and *d*_confocal_ = 240 nm, x-axis: correlation lag time τ, y-axis: line pixels i.e. space, color code: normalized G(τ) decaying from red to blue). **E)** LIESS-FCS, for each pixel of the scanned line, allows to calculate values of D_rat_ (D_rat_ = D_STED_/ D_conf_) that is obtained by the analysis of the correlation carpets in (**C)** and (**D)** which can be summarized in **F)** a frequency histogram indicating fluctuation around D_rat_ = 1 i.e. free diffusion (red line in **(E)**).

We first set out to validate LIESS-FCS using Monte Carlo simulation of freely diffusing molecules in a 2D plane. Fig. 1E depicts resulting representative values of D_rat_ for each pixel along the line, which as expected for the simulated free diffusion fluctuates around 1.0 without spatial heterogeneity and can also be displayed as a D_rat_ histogram for clarity (Fig. 1F). It is expected that the accuracy of the acquired D_rat_ values would be highly dependent on the signal-to-noise-ratio (SNR) of the measurement (a general rule for scanning-FCS measurements^21^). Note that the SNR is more impaired in the LIESS-FCS modality using alternating lasers, particularly because the total signal is split into two channels (the confocal and STED), i.e. it is halved compared to conventional sSTED-FCS recordings. As expected, the determined variability in D_rat_ values reduces (i.e. the accuracy increases) with increasing acquisition time, thus increasing amount of total signal (from 5 to 40 s, Fig. S1).

Next, we tested LIESS-FCS experimentally, and compared its performance with standard sSTED-FCS. We first used a fluorescent lipid analogue (Abberior Star Red labelled 1,2-Dipalmitoyl-sn-glycero-3-phosphoethanolamine (DPPE)) freely diffusing in a fluid supported lipid bilayer (SLB, composed of 50 % 1,2-dioleoyl-sn-glycero-3-phosphocholine (DOPC) and 50 % cholesterol). Fig. 2A and 2B show the obtained correlation carpets in STED (*d*_STED_ = 100 nm) and confocal (*d*_conf_ = 240 nm) modes, which appear very similar for conventional sSTED-FCS and LIESS-FCS, respectively. The average transit times τ_D_ obtained from fitting all correlation data of the carpets were as well similar for both approaches (Fig. 2C). Values of D_rat_ as determined from LIESS-FCS fluctuate around 1.0 without significant spatial heterogeneity as expected for free diffusion (Fig. 2D, E). As anticipated from the simulated data, the accuracy of determining D_rat_ increased with measurement time (from 10 to 40 s, Fig. S2).

**Figure 2:**
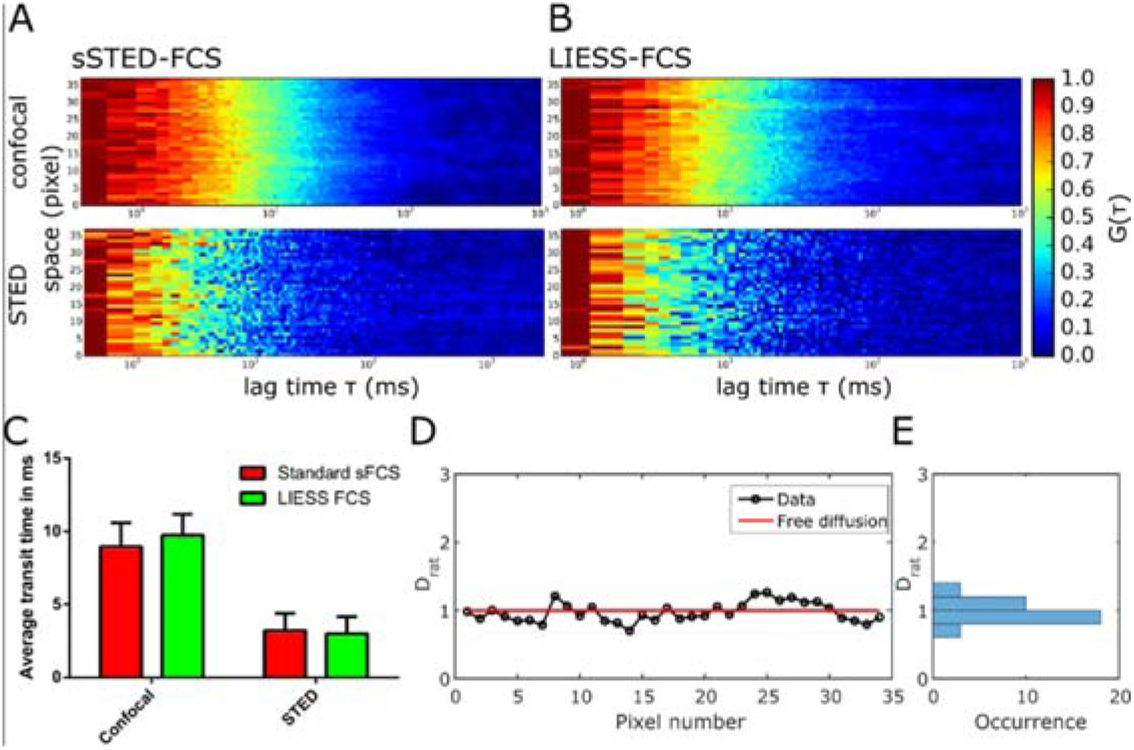
Experimental LIESS-FCS recordings of free diffusion in SLBs (Abberior Star Red labelled DPPE in DOPC/Cholesterol). **A)** Representative correlation carpets of confocal (*d*_conf_ = 240 nm, upper panel) and STED (*d*_STED_ = 100 nm, lower panel) from conventional sSTED-FCS and **B)** LIESS-FCS correlation carpets in confocal (*d*_conf_ = 240 nm, upper panel) and STED (*d*_STED_ = 100 nm, lower panel, measurement time 150 s, 1.36 µm scan). **C)** Values of transit times (average and standard deviation of the mean as error bars) determined from confocal and STED correlation carpets of the sSTED-FCS and LIESS-FCS recordings (72 curves in confocal and STED), indicating no significant difference between sSTED- and LIESS-FCS. **D)** Values of D_rat_ along the pixels of the scanned line resulting from the analysis of the LIESS-FCS correlation carpets and **E)** frequency histogram indicating fluctuations around D_rat_ = 1.0 i.e. free diffusion (red line in D).

Lipids are shown to exhibit different diffusion characteristics that is tightly linked to their function. Therefore, we next used LIESS-FCS to further investigate the diffusional characteristics of fluorescently labelled DPPE (Atto647N-labelled DPPE) and compare it with sphingomyelin (Atto647N-labelled SM) in the plasma membrane of live PtK2 cells. Previous sSTED-FCS experiments have demonstrated mainly free homogeneous diffusion for DPPE and spatially distinct spots of slowed down diffusion in the case of SM, only visible in the STED recordings^16^. However, due to the lack of simultaneous information from confocal recordings (e.g. slowed down diffusion at the same locations), this observation using sSTED-FCS could not directly be attributed to trapping interactions as reported from single-point STED-FCS measurements^13,19,20^. Fig. 3 shows representative LIESS-FCS data (correlation carpets in STED (*d*_STED_ = 100 nm) and confocal (*d*_conf_ = 240 nm) modes as well as values of D_rat_ over space) for DPPE (Fig.3A – C) and SM (Fig.3D – F). For sSTED-FCS data, the correlation carpets of the STED recordings demonstrate the appearance of spots of slowed down diffusion in the case of SM unlike DPPE (Fig.3A and 3D). The LIESS-FCS modality now allows to directly link these spots to trapped diffusion, since D_rat_ is << 1.0 at these spatial positions only (highlighted by the numbers in Fig. 3D and 3E and Fig. S3), while D_rat_ is close to 1.0 in-between (almost free diffusion as continuously detected for DPPE). Therefore, any other spatial heterogeneity showing up, such as already in the confocal correlation carpets of DPPE (Fig. 3A and 3B arrows in the correlation carpets and D_rat_ plot), are still characterized by free diffusion, i.e. they do not relate to trapping interactions despite the obvious heterogeneity. A possible cause for such heterogeneity may be the uneven plasma membrane topology involving curvatures^16^.

**Figure 3:**
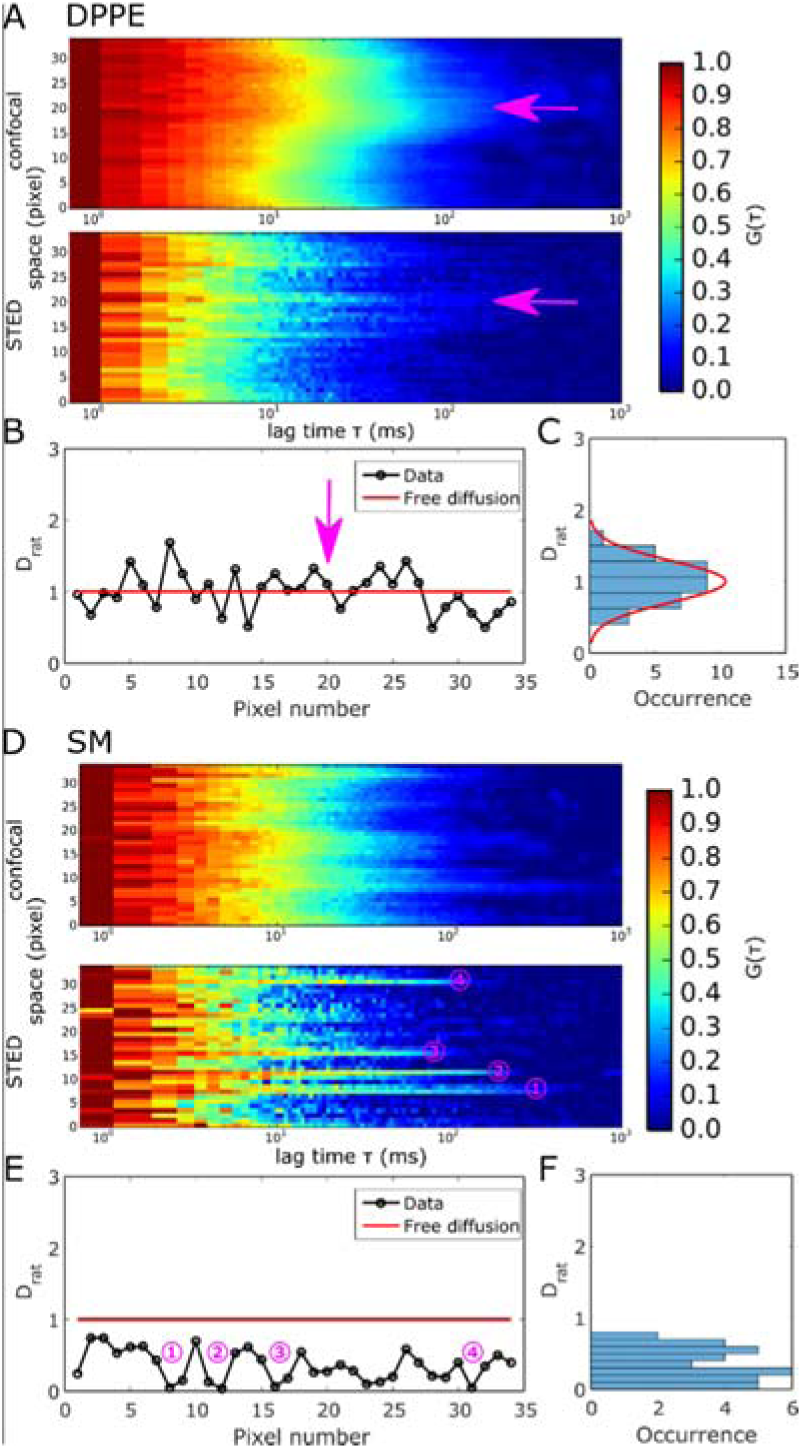
Experimental LIESS-FCS recordings for Atto647N-labelled DPPE (**A - C**) and Atto647N-labelled SM (**D - F**) in the plasma membrane of live PtK2 cells. **A)** Representative correlation carpets of simultaneous confocal (*d*_conf_ = 240 nm, upper panels) and STED (*d*_STED_ = 100 nm, lower panels) recordings for DPPE (measurement time 120 s, 1.36 µm scan). **B)** Values of D_rat_ resulting from the correlation carpet analysis and **C)** frequency histogram indicating fluctuation around D_rat_ = 1 i.e. free diffusion for DPPE. The arrows indicate an exemplary area where heterogeneity is still characterized as free diffusion. **D)** Representative correlation carpets of confocal (*d*_conf_ = 240 nm, upper panels) and STED (*d*_STED_ = 100 nm, lower panels) recordings for SM (measurement time 45 s, 1.36 µm scan). **E)** Values of D_rat_ resulting from the correlation carpet analysis and **F)** frequency histogram indicating trapping sites (D_rat_ << 1). Numbers in (**D**) and **(E)** show the exact same trapping sites.

To better understand the temporal organisation of the depicted trapping sites for SM, we split the longer LIESS-FCS data into 30 s measurements. The respective correlation carpets as well as spatially-resolved values of D_rat_ reveal a transient character of the sites, i.e. trapping sites disappeared and new ones appeared (Fig. 4) (either due to some sort of molecular assembly/disassembly or diffusion) which is in accordance with the transient character of the spots of slowed-down diffusion observed in the previous sSTED-FCS recordings^16^. Since still dominating the 30 s recordings, the trapping sites have to be stable for at least a few seconds. This transient character brings up an issue of the duration of a LIESS-FCS measurement, since too long acquisition times (which are definitely favourable for improved statistical accuracy, compare Fig. S1 and S2) may average over the appearance and disappearance of the trapping sites. This is exemplified in Fig. S4 which shows the correlation carpet and spatially resolved values of D_rat_ for different acquisition time windows (0-10s, 0-20s … 0-100s) of the same LIESS-FCS recording. It becomes obvious that too short acquisition times (10 s) result in noisy data (as highlighted by spikes towards values of D_rat_ >> 1.0), while too long acquisition times (>40 s) average over appearing and disappearing trapping sites resulting in rather spatially homogeneous values of D_rat_ < 1.0 (as over time almost every pixel along the scanned line has experienced a trapping site). Note, that the 100 s recording for DPPE still resulted in continuous values of D_rat_ = 1.0, precluding the appearance of dominant trapping sites for this lipid analogue. Finally, relating our current LIESS-FCS data to the previous point and scanning STED-FCS data^13,16,19,20^, we can conclude that the trapping sites are smaller than 80 nm in size, transient in the second-time range, and that certain lipids such as SM transiently (over a few ms) interact with entities in these hot-spots.

**Figure 4:**
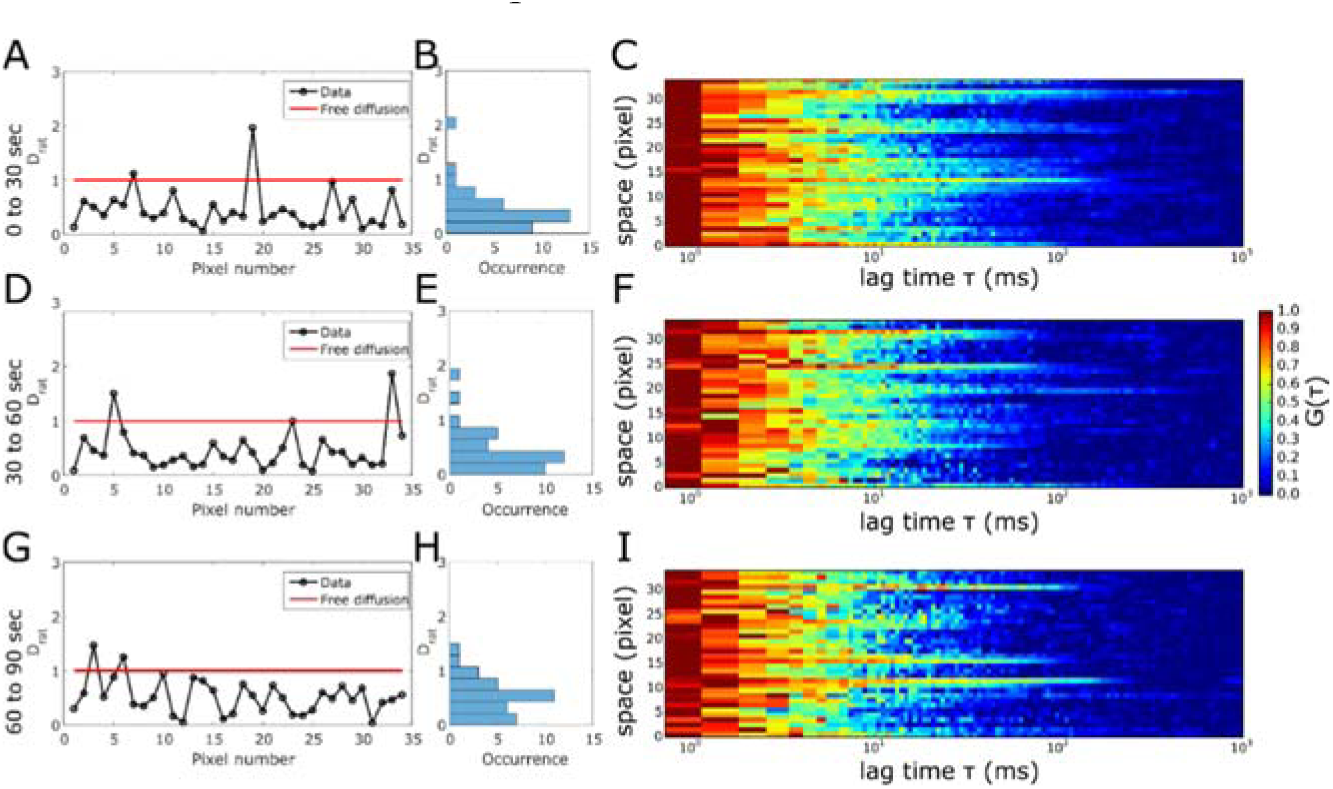
Transient nature of the trapping hot-spots; temporal cropping of the experimental LIESS-FCS recordings for Atto647N-labelled SM in the plasma membrane of live PtK2 cells. Measurement times as marked: **A-C)** 30-60 s, **D-F)** 30-60 s, **G-I)** 60-90 s. **A**, **D**, **G)** D_rat_ values for all the pixels of the scanned lines, **B**, **E**, **H)** frequency histograms and **C**, **F**, **I)** representative correlation carpets of STED recordings (*d*_STED_ = 100 nm) indicating trapping sites (D_rat_ << 1) and fluctuations in-between the subsequent recording.

Finally, we employed LIESS-FCS to investigate the diffusional behaviour of GPI-anchored proteins (GPI-APs) which play a major role in various cellular signalling pathways. Their spatio-temporal organisation is quite controversial^22–24^ and can now be tackled with our technique. Fig. 5 depicts representative LIESS-FCS data for a GPI-AP in the cellular plasma membrane of live PtK2 cells. We utilized a GPI-anchored SNAP-tag (GPI-SNAP) as a representative GPI-AP^20^. Fig. 5A shows a confocal image of the basal plasma membrane of a live PtK2 cell transfected with a GPI-SNAP (labelled with the dye Abberior STAR Red), indicating an almost homogeneous distribution with only a few bright spots. Such bright spots were observed before for such GPI-APs^20^ and they were associated with bright and immobile GPI clusters or assemblies at the close-vicinity of the plasma membrane. Crossing of these isolated bright GPI-AP clusters during beam-scanning should in principle be avoided in scanning-FCS measurements, since such immobile features usually bias the data due to photobleaching, appearing as correlation curves with prolonged decay times^21^. Such long decays also appear in some locations of the representative correlation carpet shown in Fig. 5B. Yet, as the photobleaching-based bias affects both the confocal and STED correlation carpets at the same position, these events can straightforwardly be assigned to a photobleaching artefact (while they may accidentally be considered as trapping sites in standard sSTED-FCS recordings (Fig. S5A)). Concerning the mobile pool, the diffusion modes of GPI-SNAP turned out to be quite heterogeneous. As shown in the representative data of Fig. 5C and 5D and Fig. S5B, we observed values of D_rat_ ranging from <<1 (trapping) over 1 (free) to >1 (hop). This is confirmed by the broad histogram of D_rat_ values gathered from LIESS-FCS measurements on 5 different cells, tailing into values D_rat_ > 1 (Fig. 5E), and its peak value of D_rat_ = 0.6 highlights a dominant trapping diffusion character. We have to note that this heterogeneity came apparent despite a rather long measurement time of 70 s which excludes the possibility of noise-related heterogeneity.

**Figure 5:**
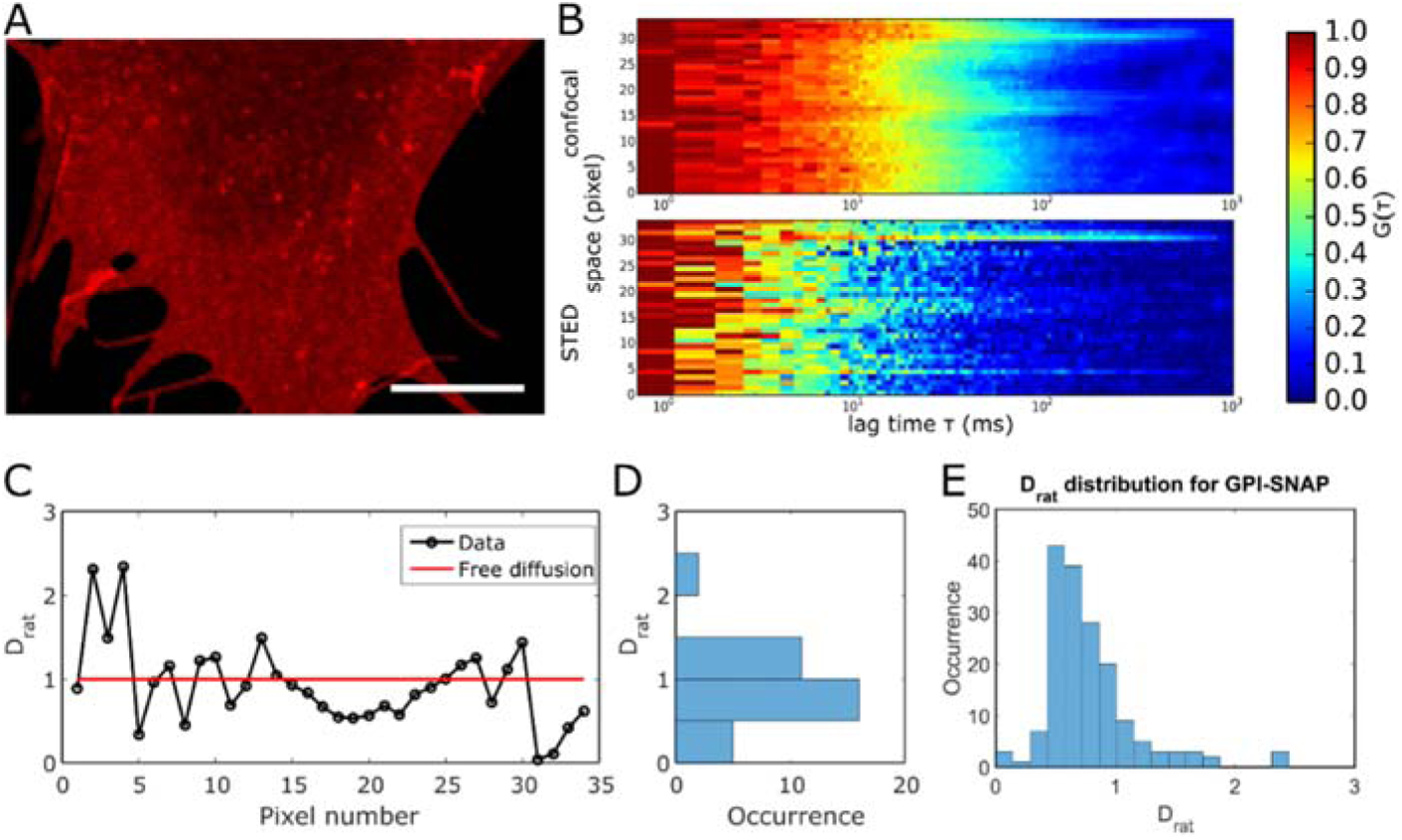
Experimental LIESS-FCS recordings of the fluorescently tagged (Abberior STAR Red) GPI-SNAP protein in the plasma membrane of live PtK2 cells. **A)** Representative confocal image of a portion of the cellular membrane indicting homogeneous distribution with rare bright clusters. Scale bar: 10 µm. **B)** Representative correlation carpet of the simultaneous confocal (*d*_conf_ = 240 nm) and STED recordings (*d*_STED_ = 100 nm, measurement time 70 s, 1.36 µm scan). **C)** Values of D_rat_ resulting from the analysis of the correlation carpet with **D)** frequency histogram indicating large fluctuation of D_rat_. **E)** Histogram of values of D_rat_ obtained from 5 different line scans on 5 different cells with a peak at D_rat_ = 0.6 and a broad distribution with values ranging from Drat << 1 (trapping), D_rat_ = 0 (free) to D_rat_ > 1 (hop), confirming the strong variation in diffusion modes.

Our data demonstrate the capability of LIESS-FCS to directly observe spatial heterogeneity in molecular diffusion behaviour (such as spatially distinct sites of trapping, hop or free diffusion). The strength of LIESS-FCS results from the simultaneous acquisition of confocal and STED-FCS data at different spatial positions. Unfortunately, this comes with the price of a low SNR, which demands rather moderate acquisition times of 30-100 s and moderately reduced observation spots *d* ≈ 100 nm. A remedy may be the use of dyes with even further increased fluorescence yield or the use of time-gated detection schemes^17^ or phasor-plot analysis^25^. Nevertheless, LIESS-FCS provides an unique tool for investigating lateral organisation of cellular membranes on variable length scales accounting for bias due to biological heterogeneity or photobleaching artefacts, possibly answering long-standing questions of functional membrane heterogeneity^1,2^.

## Materials and Methods

### SLB preparation

SLBs were prepared by spin coating lipid mixtures as described previously for pure DOPC bilayers^20^. A solution with a total concentration of 1 mg/mL of 1,2-dioleoyl-sn-glycero-3-phosphocholine (DOPC, Avanti Polar Lipids) and cholesterol (Avanti Polar Lipids) at a molar ratio of 0.5 in chloroform:methanol (1:2) was doped with 1:2000 fluorescent lipid (Abberior STAR Red DPPE, Abberior) and was spin coated at 3200 rpm onto a clean 25 mm round microscope cover slip. The SLB was formed after hydrating the lipid film with SLB buffer (150 mM Nacl, 10 mM HEPES). The SLB was stable for hours. Prior to coating, the microscope cover slips were cleaned by etching with piranha acid. Fresh cover slips were stored for no longer than one week.

### PtK2 cell handling and labelling

PtK2 cells were kept in DMEM (Sigma Aldrich) supplemented with 1 mM L-Glutamin (Sigma Aldrich) and 15 % FBS (Sigma Aldrich). For experiments, cells were seeded onto 25 mm round microscope cover slips kept in 35 mm petri dishes. After allowing the cells to grow for 24-48 hours and reaching a confluency of roughly 75 %, cells were ready for experiments. After washing with L15 (Sigma Aldrich) cells were labelled for 15 minutes with fluorescently lipid analogues (Atto647N-DPPE and Atto647N-SM, Atto-Tec) at a concentration of 0.4 µg/mL and subsequently washed with L15. Including labelling the cells were kept for not longer than 1 hour at room temperature. Measurements were performed at room temperature to prevent internalisation of the lipid analogues. Transfection of PtK2 cells was performed using Lipofectamine 3000 (Life Technology) according to the manufacture’s protocol. The medium was exchanged 3 hours after transfection. GPI-SNAP (kind gift from the lab of Stefan Hell) was labelled with the non-membrane permeable SNAP ligand Abberior STAR Red for 45 minutes in full medium at 37 °C. The cells were washed two times for 15 minutes with full medium at 37 °C and subsequent measurements were performed in L15 for not longer than 1 hour at room temperature.

### Data acquisition and fitting

All scanning STED-FCS and LIESS-FCS data were acquired at a customised Abberior STED/Resolft microscope as previously described^20^. The data acquisition was controlled with Abberior’s Imspector software. The scanner was optimised for sFCS. Standard sFCS data were obtained from an x-t scan. Measurement times were between 30 seconds and 180 seconds. For LIESS-FCS, we made use of the Line Step function alternating the excitation between confocal and STED mode between every other scanned line, and the intensity data for confocal and STED modes were sorted into two independent channels. Typically sFCS acquisition was performed using an orbital scan with a pixel dwell time of 10 µs and scanning frequencies of about 3 kHz. The pixel size was kept to 40 nm resulting in an orbit with a diameter of roughly 1.5 µm. Control sFCS measurements were performed with a frequency of roughly 1.5 kHz, a pixel dwell time of 10 µs and an orbit with a diameter of 3 µm.

Confocal and STED microscopy performances were checked using 20 nm Crimson beads on a daily basis. The diameter *d*_*STED*_ of the observation spots in the STED mode were deduced from measurements of the freely-diffusing fluorescent lipid analogue Abberior Star Red DPPE. *d*_*STED*_ was calculated from the diameter of the confocal observation spot *d*_conf_ (as determined from Crimson bead measurements) and the transit times in confocal (*τ*_*D,confocal*_) and STED (*τ*_*D,STED*_) mode:

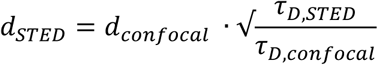

We usually employed *d*_*STED*_ ≈ 100 nm in our LIESS-FCS measurements; smaller diameters as realized in previous single-point and scanning STED-FCS experiments resulted in too noisy correlation data.

For analysis, the x-t intensity carpets (temporal fluorescence intensity data for each pixel) were correlated and subsequently fitted using the conventional model for 2D-diffusion in a plane:

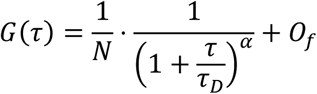

in the FoCuS-scan software^21^ (https://github.com/dwaithe/FCS_scanning_correlator) with *N* as the average number of molecules in focus τ_*D*_ as transit time, *α* as anomaly factor and *O*_*f*_ as offset. To remove immobile components the first 10 to 20 seconds were cropped off from all measurements. Additionally, the first pixel of the line was cropped out. In some cases, especially for cell measurements, photobleaching correction was applied (fitting the total intensity data over time with a mono-exponential decay for SM or averaging over 15s-time intervals for DPPE). Subsequently, the data were fitted with the single component diffusion model. The anomaly factor α was fixed to 1 for the simulation and SLB data but was left free-floating between values of 0.8 -1.05 for cellular data^13^. To obtain stable fits, the data were bootstrapped 20 to 40 times^21^. From the obtained transit times in confocal and STED the apparent diffusion coefficient D_app_ was calculated according to:

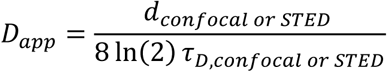

The values of D_rat_ = D_app_(STED)/D_app_(confocal) over space x were generated using a custom written Matlab script.

### Simulations of free diffusion

To validate our approach we performed Monte Carlo simulations using the nanosimpy library in Python (https://github.com/dwaithe/nanosimpy) as described previously^21^. Freely moving particles were simulated in a box of 2 µm times 8 µm. In case of a molecule hitting the edges of the box it was wrapped around to appear on the opposite side (diffusion on a sphere). The sFCS line was placed in the centre of the box with its ends at least 1 µm away from the boundaries. Molecules were passed through a Gaussian shaped observation spot as appropriate. To mimic the LIESS-FCS measurements, data were obtained alternating between confocal and STED observation spots mimicked by a Gaussian with a FWHM of 240 nm and 100 nm, respectively. The resulting intensity carpets were saved as. tiff files, correlated and analysed as described above for the experimental data.

## Acknowledgements

We thank the Wolfson Imaging Centre Oxford and the Micron Advanced Bioimaging Unit (Wellcome Trust Strategic Award 091911) for providing microscope facility and financial support. ES is funded by EMBO Long term and Marie Skłodowska-Curie Intra-European (MEMBRANE DYNAMICS) postdoctoral fellowships. We acknowledge funding by the Wolfson Foundation, the Medical Research Council (MRC, grant number MC_UU_12010/unit programmes G0902418 and MC_UU_12025), MRC/BBSRC/EPSRC (grant number MR/K01577X/1), the Wellcome Trust (grant ref 104924/14/Z/14), the Deutsche Forschungsgemeinschaft (Research unit 1905 “Structure and function of the peroxisomal translocon”), Newton-Katip Celebi Institutional Links grant (352333122) and Oxford-internal funds (John Fell Fund and EPA Cephalosporin Fund). J.B.S. acknowledges financial support by the Marie Curie Career Integration Grant “NanodynacTCELLvation” PCIG13-GA-2013-618914.

