## Supplementary Materials for "Nanoscale spatio-temporal diffusion modes measured by simultaneous confocal and STED imaging"

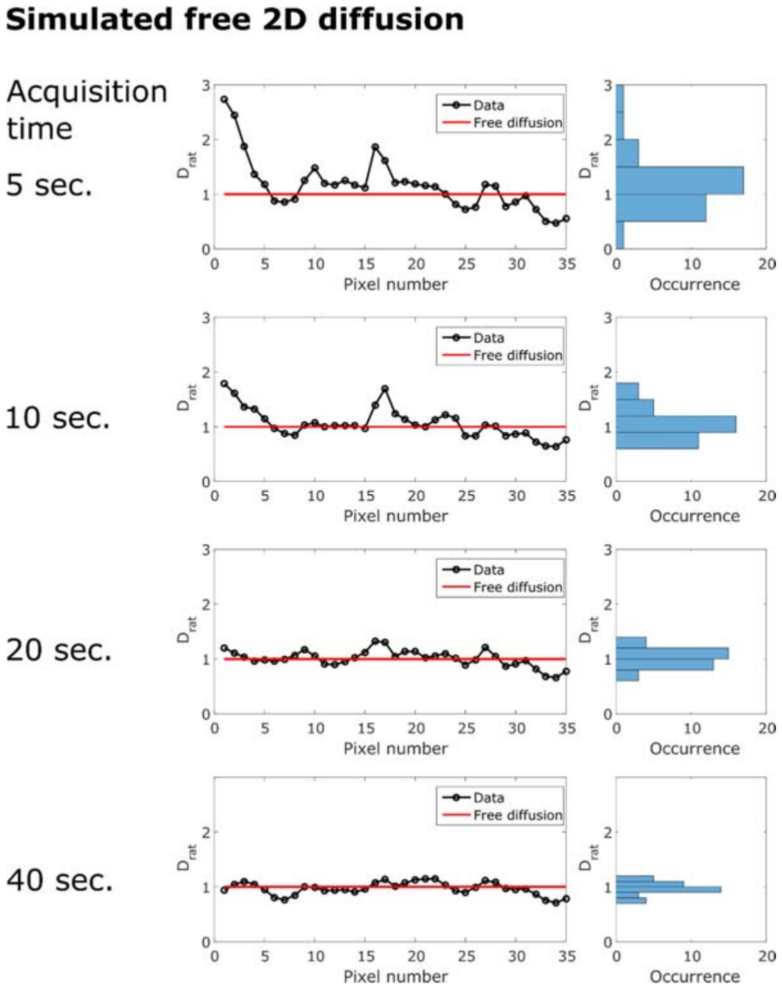


**Figure S1**: Monte Carlo Simulations of freely diffusing molecules measured with LIESS-FCS and sampled with different acquisition times (1.36 µm scan). The $D_{rat}$ values over space (left) and the $D_{rat}$ histograms (right) fluctuate around $D_{rat}$ = 1.0 improving in signal-to-noise with increasing acquisition times as here demonstrated by varying the bin edges towards the minimum and maximum values of $D_{rat}$ in the histograms on the right.

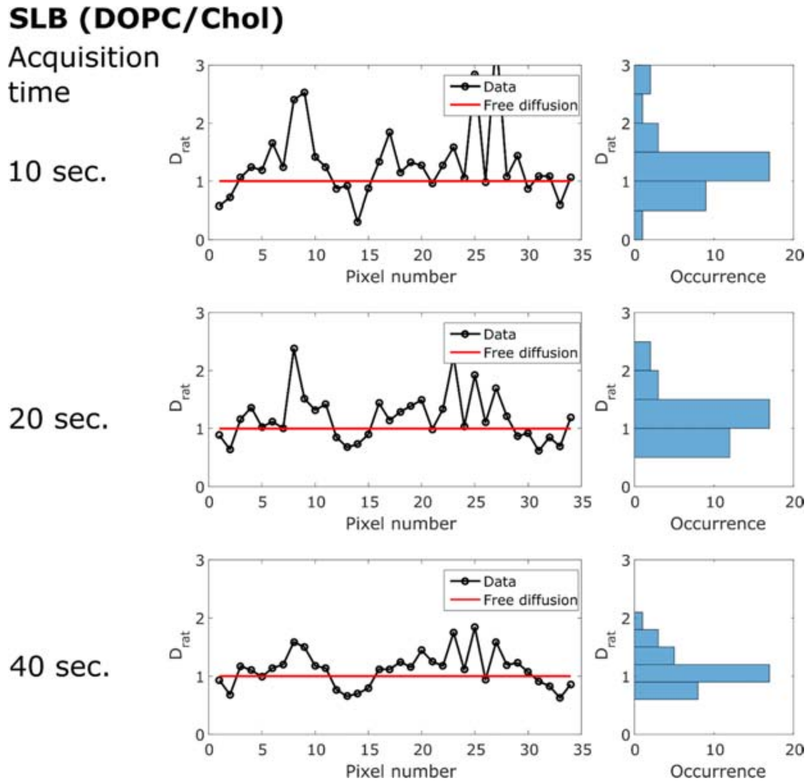


**Figure S2**: LIESS-FCS measurement of freely diffusing lipid (Abberior STAR Red DPPE) within a SLB composed of DOPC/Cholesterol (50 mol% each) cropped to different acquisition times (1.36 µm scan). The $D_{rat}$ values over space (left) and the $D_{rat}$ histograms (right) fluctuate around $D_{rat}$ = 1.0 improving in signal-to-noise with increasing acquisition time.

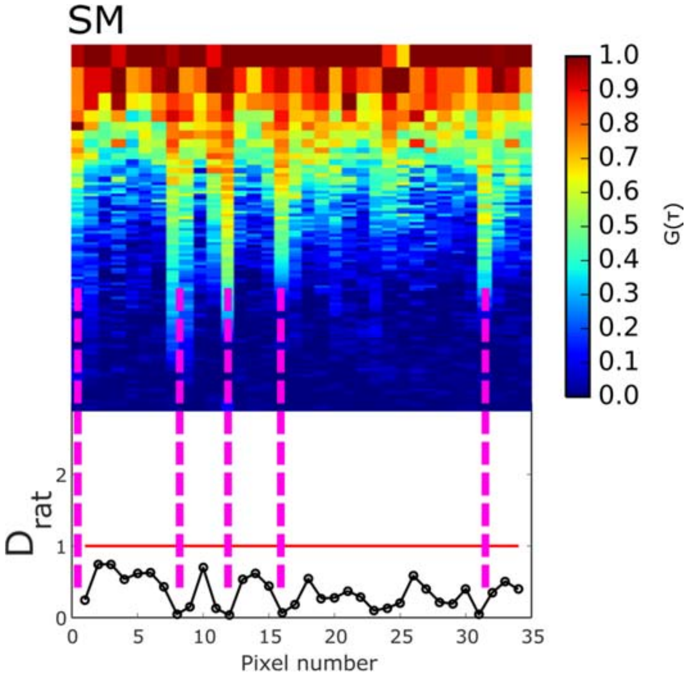


**Figure S3:** Experimental LIESS-FCS recordings of diffusion of the fluorescent SM. The trapping sites in the correlation carpet (top) align with particularly low values of $D_{rat}$ in the plots over space (bottom) as highlighted by the dashed lines (measurement time of 60 seconds, 1.36 μm scan).

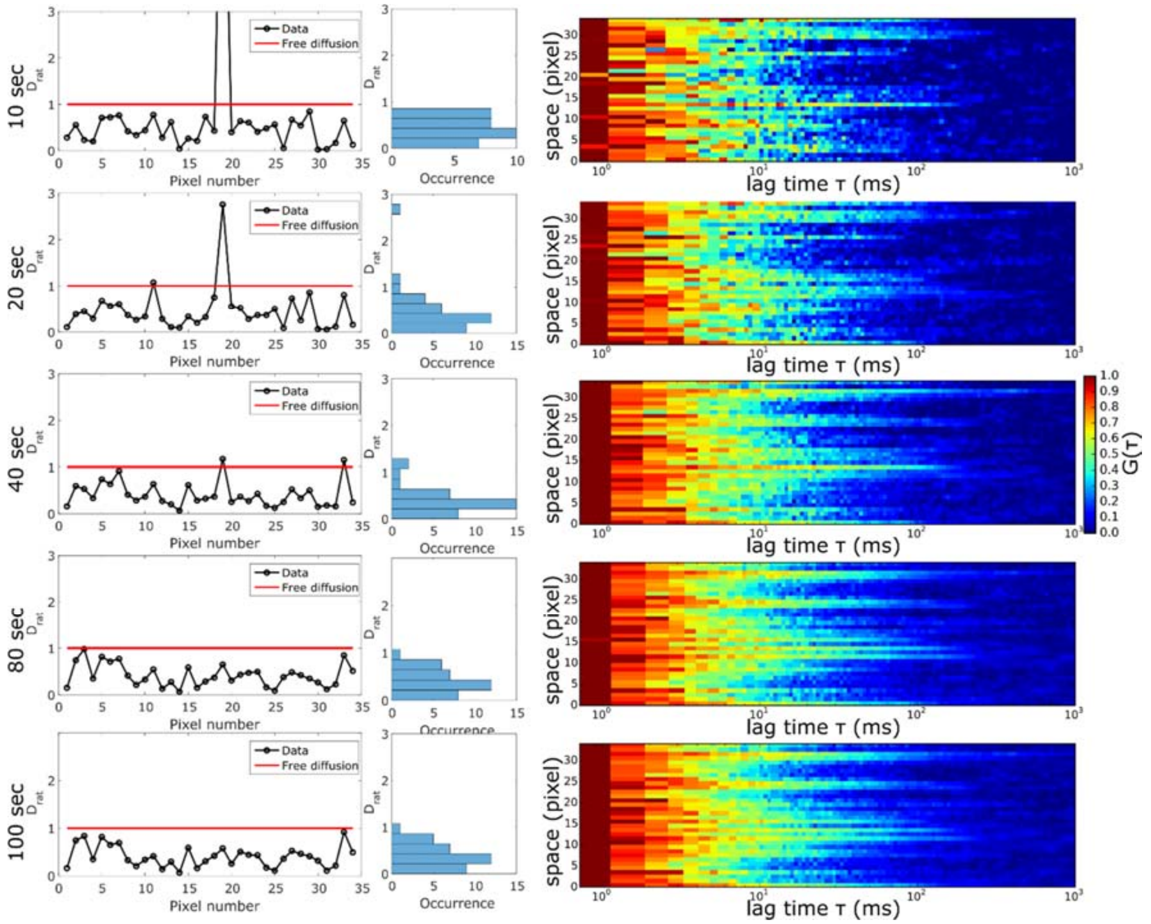


**Figure S4**: Experimental LIESS-FCS recording of fluorescent SM analogue in live PtK2 cells cropped to different accumulation times. The effect of measurement time and averaging on the trapping effect is exemplified on the $D_{rat}$ over space plots (left) and the correlation carpets in STED (right). With longer acquisition time $D_{rat}$ notably shifts towards 0 whereas noise dominates measurements with short acquisition times. Sufficient signal to noise seems to be obtained at a measurement time of about 40 seconds.

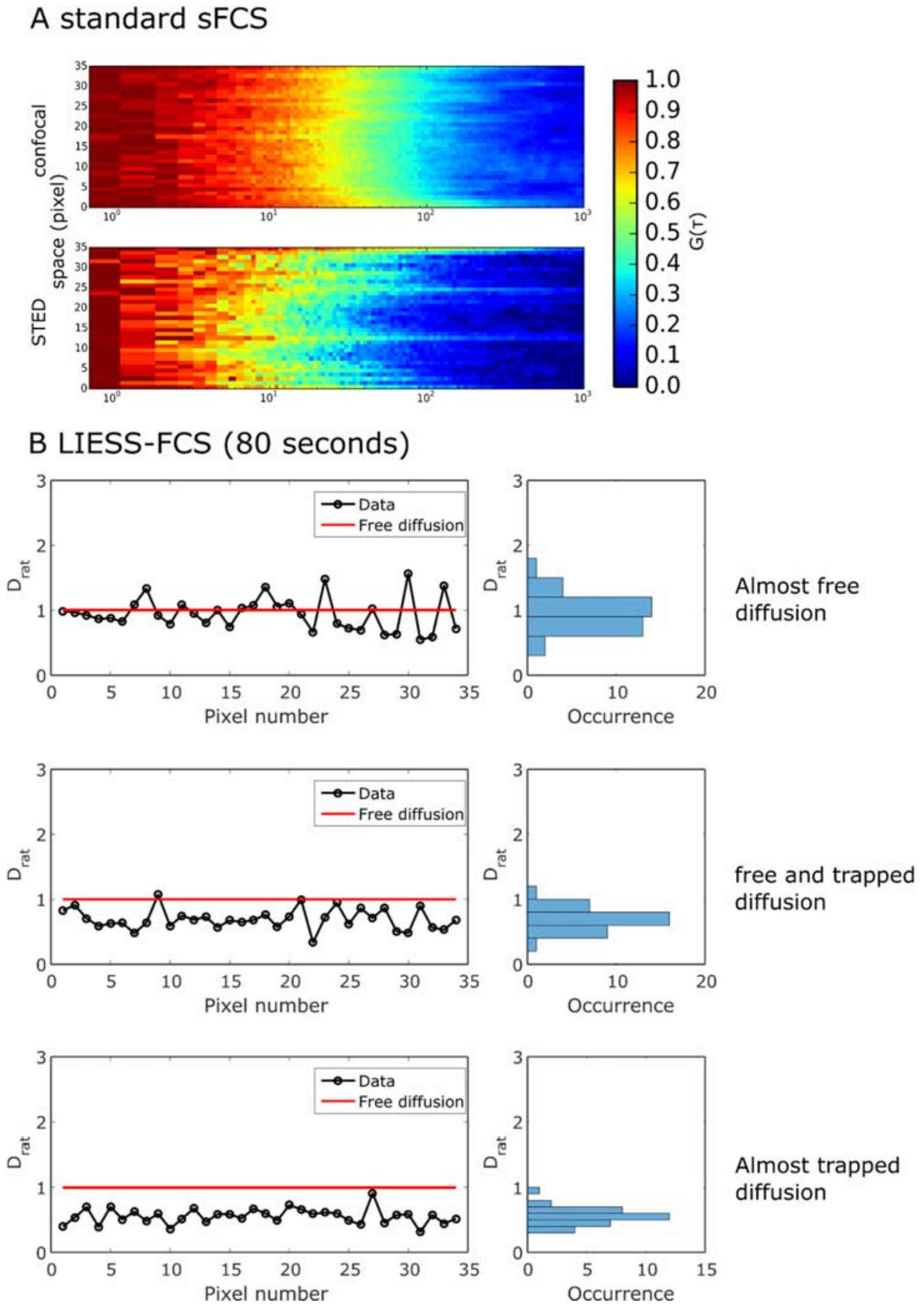


**Figure S5**: Experimental standard sFCS and LIESS-FCS recordings of fluorescently tagged (Abberior STAR RED) GPI-SNAP. **A)** Standard confocal and STED sFCS correlation carpets (measurement time 80 s, 1.36 µm scan) with strong heterogeneities apparent in STED. **B)** LIESS-FCS recordings of three different cells (measurement time 80 s, 1.36 µm scan). The $D_{rat}$ plots over space (left) and the $D_{rat}$ histograms (right) display free diffusion, trapped diffusion and a mixture.
